# Biobtree: A tool to search, map and visualize bioinformatics identifiers and special keywords

**DOI:** 10.1101/520841

**Authors:** Tamer Gur

## Abstract

Due to their nature, bioinformatics datasets are often closely related to each other. For this reason, search, mapping and visualization of these relations are often performed manual or programmatically via identifiers or special keywords such as gene symbols. Although various tools exist for these situations, the growing volume of bioinformatics datasets, emerging new software tools and approaches motivates new solutions. To provide a new tool for these current cases, I present the Biobtree bioinformatics tool. Biobtree effectively fetches and indexes identifiers and special keywords with their related identifiers from supported datasets, optionally with user pre-defined datasets and provides a web interface, web services and direct B+ tree data structure–based single uniform database output. Biobtree can handle billions of identifiers and runs via a single executable file with no installation and dependency required. It also aims to provide a relatively small codebase for easy maintenance, addition of new features and extension to larger datasets. Biobtree is available to download at https://www.github.com/tamerh/biobtree.

## Introduction

Bioinformatics datasets often consist of entries, where each entry is represented by unique identifier. Depending on the dataset, each entry contains various types of information such as biological function, chemical structure or literature reference etc. In addition, entries often contain cross-referencing information to other dataset entries via identifiers. Let’s take as an example entry the proto-oncogene vav protein in humans, which is encoded by the *VAV1* gene. If we display this protein on the UniProt <u>website,</u> we see cross references to many other databases. These cross references represent relations of databases with each other. Various tools exist to deal with such data; however, the growing volume of bioinformatics datasets, emerging new software tools and analysis approaches motivates new solutions. Biobtree presented herein is capable of improved and rapid processing of large numbers of unique identifiers of entries and related identifiers that are specified via cross-reference data.

In some datasets, in addition to unique identifiers there is information that is strongly related to entries but not necessarily unique for each entry. Species names or UniProt secondary accessions are example of this type. Information that is strongly related to the entries but not necessarily unique is a second data source for Biobtree. In Biobtree, these types are called special keywords and each of these can be related to multiple entries among the datasets.

Biobtree retrieves all these identifiers, related identifiers and special keywords from various bioinformatics resources and stores it in a single database. The data resources currently used are ChEBI [1], HGNC [2], HMDB [3], InterPro [4], Europe PMC [5] and UniProt [6]. Figure 1 shows details of these datasets.

**Figure 1.**
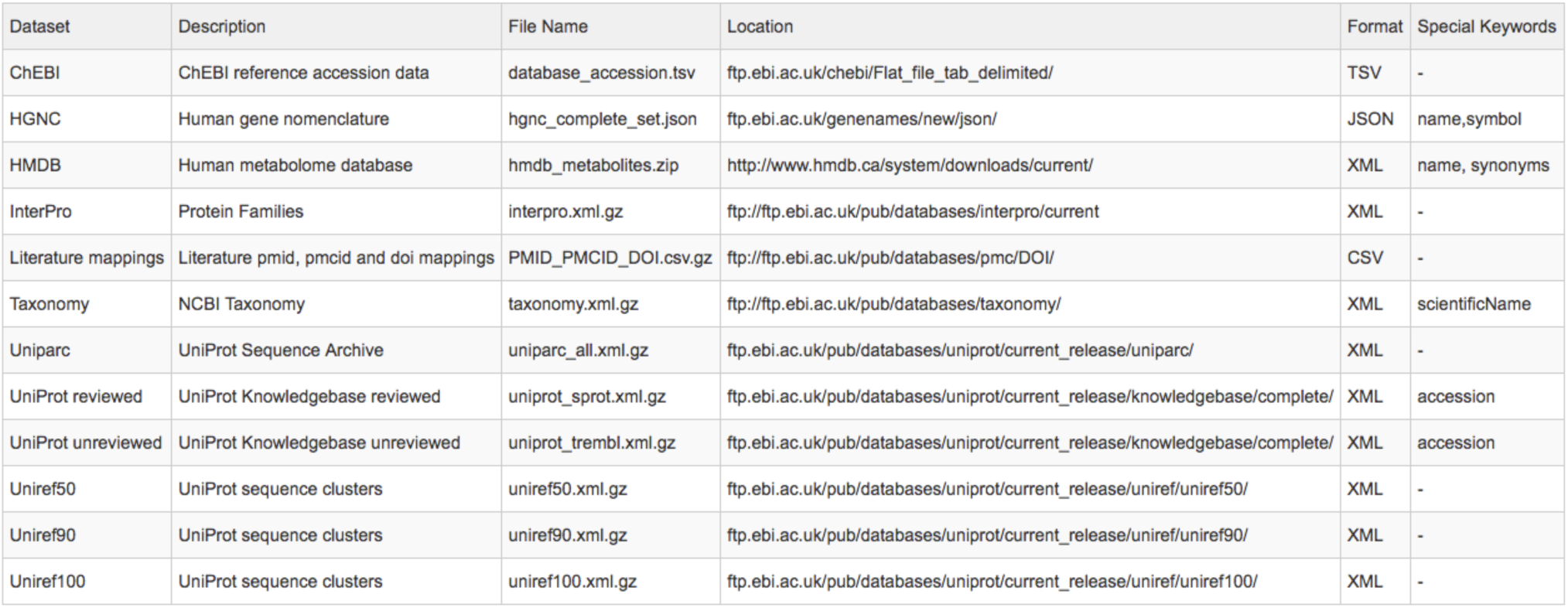
List of datasets.

Based on stored data, Biobtree provides search, map and visualization functionalities via provided web services or a web interface. For instance, all the UniProt proteins entries belonging to a gene name, or, all Ensembl [7] genome transcripts identifiers and ENA [8] sequence identifiers that map to a protein identifier can be accessed. These relations, determined via identifiers, are stored bidirectionally so all actions can also be done in the opposite way.

Biobtree is managed from a single executable file for each major operating system without requiring any installation or compilation. It consists of three main phases, which will be explained in later sections. These phases are named *update, generate* and *web* and are controlled by a Biobtree command line interface (CLI). As a database, Biobtree uses a B+ tree data structure–based <u>LMDB</u> key value store. LMDB provides fast batch inserts and reads and allows effective operation on a large number of records. LMDB is embedded into Biobtree’s executable binary code so it does not require a separate installation.

## Update Phase

The purpose of the update phase is to retrieve dataset identifiers and special keywords from remote servers or a local disk and produce files that contains identifiers and special keywords with their referred identifiers as keys and values in a sorted order. It is essential that the produced files are sorted to make fast batch inserts to LMDB database in the next phase. The updating phase is started via the Biobtree CLI with the update command. For example, the following command starts the update phase for the hgnc dataset.

biobtree --d hgnc update

Updating reads selected datasets as a stream and saves Biobtree-related data in a series of files. An advantage of reading dataset as streams is that it does not require fully downloading the dataset to the local disk. Datasets can have different formats like XML, JSON, TSV or CSV. Biobtree has specialized parsers for each dataset and parses them to produce its output files. When Biobtree runs the first time it retrieves its configuration, license and web interface files from the source code repository. Configuration files contain Biobtree runtime settings and dataset definitions.

### Integrate user dataset

User data can be integrated to Biobtree. This feature creates an alternative for data providers to serve their data. Data should be gzipped and in an xml format compliant with UniProt xml schema <u>definition</u>. After the file path of the data is configured in a Biobtree configuration file, updating starts similarly:

biobtree --d my_data update

### Updating in multiple computers

Biobtree supports executing the update phase over multiple computers. This is useful when it is necessary to use multiple computer processors at the same time such as with large datasets. The following two commands can be run on different computers with additional idx parameter to guarantee that the produced files have unique names.

biobtree --d uniparc –idx 1 update

biobtree --d uniref50 –idx 2 update

Although Biobtree supports the updating phase occurring over multiple computers, for the next phase all the produced files have to be in a single location.

## Generate Phase

The purpose of this phase is to merge all the files produced in the update phase by keeping the sorted order and generate the final key and values in the generated LMDB database. Keys consist of identifiers and special keywords and values are identifiers that are referred to by these keys with their dataset information. If the values size for each key are above a certain threshold they are saved in pages. The following command starts the generate phase.

biobtree generate

The generated database output is used in next web phase but it can be also used directly. Example source codes for using the database directly can be found in the project github page.

## Web Phase

The purpose of this phase is to provide web services and a web interface via the produced output of the generate phase. The following command starts web phase

biobtree web

With this command the Biobtree web server is started instantly, serving the REST, gRPC and web interface services.

### REST service

To make queries in the produced database, Biobtree provides RESTful endpoints with json-formatted responses. For example, each dataset has a unique identifier and other meta information like name, url template, etc. Dataset-unique identifiers are used in all the services to distinguish the dataset. These meta information is retrieved via the following endpoint:

http://localhost:8888/ws/meta

The following are used to query single or multiple identifiers or special keywords:

http://localhost:8888/ws/?idlist=vav_human

http://localhost:8888/ws/?idlist=vav_human,tpi1,brca2

To make a paging query for a certain identifier:

http://localhost:8888/ws/?id=vav_human&dataset=1ypage=1

To make a filtering query based on a dataset:

http://localhost:8888/ws/?id=vav_human&dataset=1&filters=102

To make a paging query with active filtering:

http://localhost:8888/ws/?id=vav_human&dataset=1&filters=102&page=1

### gRPC service

RESTful services are often json-based and are very convenient in json-based applications. But for non–json-based applications, it requires an extra process of serialization and deserialization. To address this, Biobtree provides a gRPC service with same functionality as its RESTful service. Sample codes for using gRPC in different languages can be found on the project github page. The following is snippet of Biobtree gRPC service definitions:

~~~
service BiobtreeService {
rpc Get     (BiobtreeGetRequest)     returns (BiobtreeGetResponse);
rpc GetPage (BiobtreeGetPageRequest) returns (BiobtreeGetPageResponse);
rpc Filter  (BiobtreeFilterRequest)  returns (BiobtreeFilterResponse);
rpc Meta    (BiobtreeMetaRequest)    returns (BiobtreeMetaResponse);
}
~~~

### Web interface

The Web interface allow user to visualize the produced database via the RESTful service. Once the web phase is started it is accessed via the browser from the following address:

http://localhost:8888/ui

The Web interface provides searching of multiple identifiers and special keywords, visualizing and filtering results, and executing bulk queries. On the result page for each result, it provides a url to access the main website where the data are originally produced. Figures 2 and 3 show the main and result page of the web interface.

**Figure 2.**
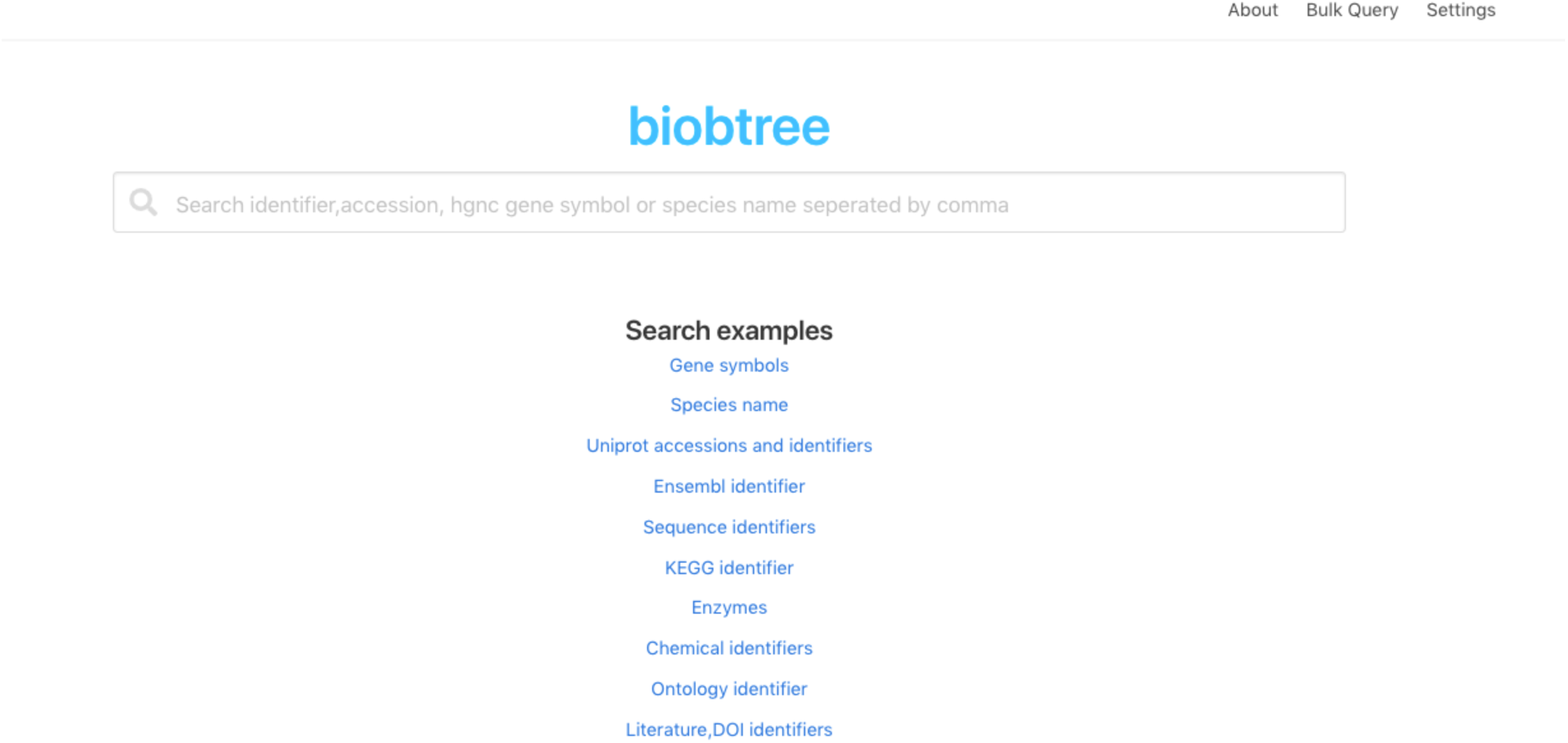
Web interface main page.

**Figure 3.**
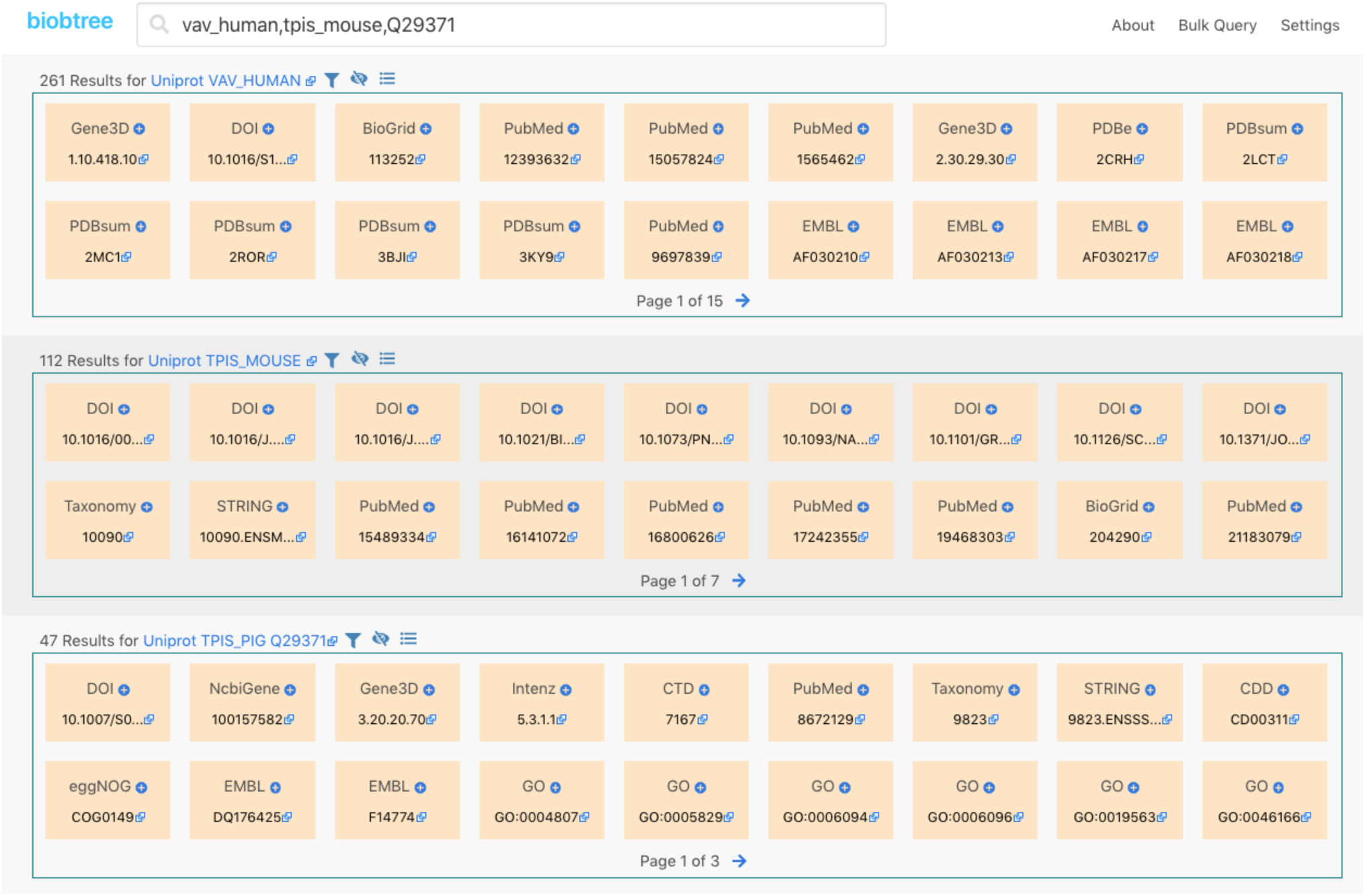
Web interface result page.

## Benchmarks

Software benchmarks should often be taken with a grain of salt, especially where the input data is large, because benchmarks results can be affected by many factors like the operating system, storage, network speed, input data, application parameters etc. Considering these, the purpose of Biobtree benchmarks is mainly to show overall capabilities and resource usage of Biobtree. Figure 4 shows the benchmark details and results. Benchmarks were primarily computed at <u>digitalocean</u> London datacentres using their cpu optimized droplets with block storage volumes. Hundred thousand sample query which used in the benchmarks can be found project github page.

**Figure 4.**
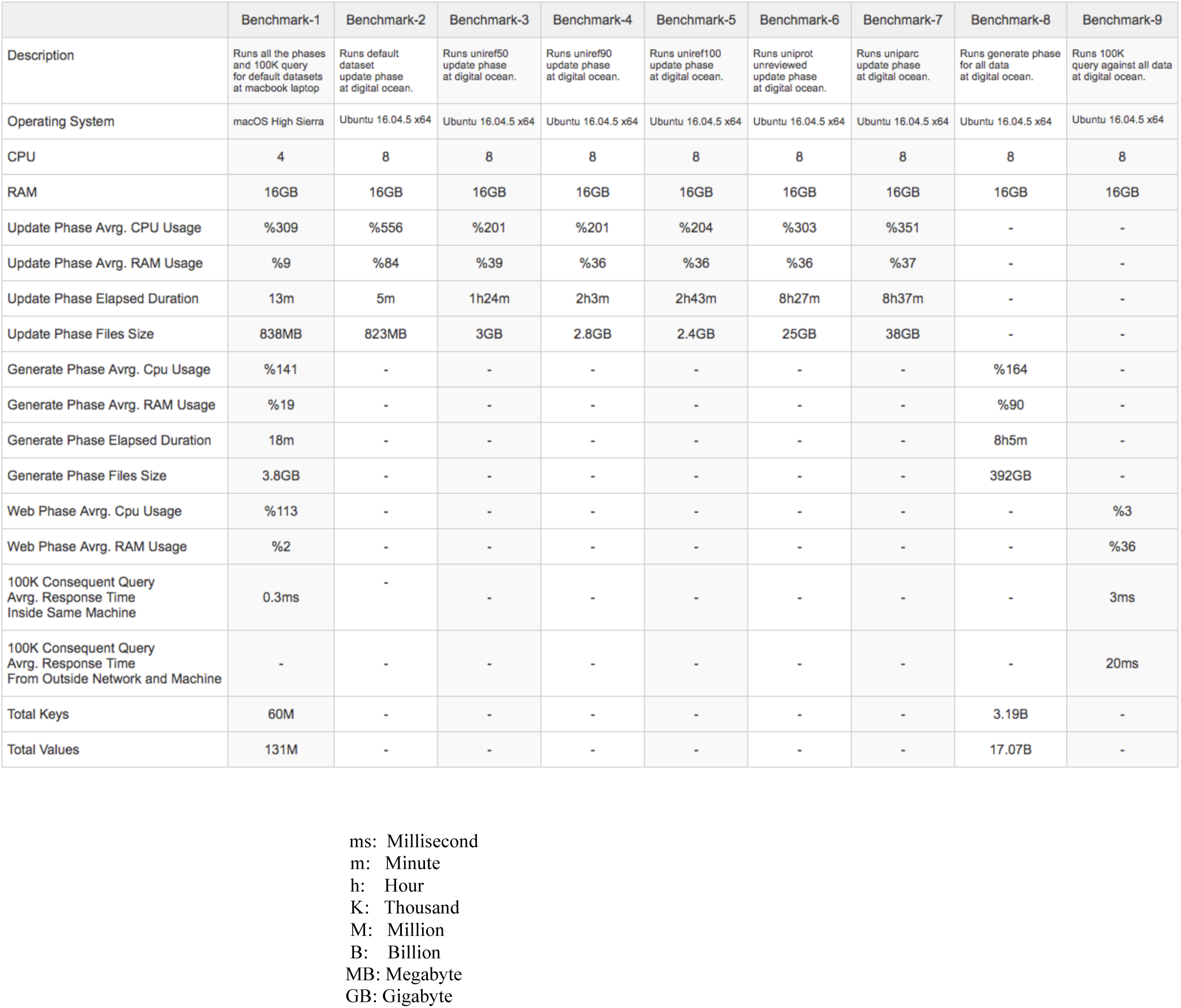
Benchmarks results.

## Discussion

The benchmarks show that Biobtree has produced LMDB database output in relatively acceptable times. Let’s discuss how Biobtree behaves if UniProt provides tens of times larger data. Clearly, more disk space would have been needed.

If enough disk space is provided, the next obstacle would have happened during the update phase, because currently UniProt provides a single gzip compressed file for each dataset and Biobtree reads each file as a stream from the beginning to end. Gzip does not allow a random access to a file unless there are checkpoints defined. This characteristic of gzip prevents the processing single large file in a split manner and utilizes more computing resources if available. Two solutions can address the issue. The first could be if UniProt would allow its datasets to be capable of parallel processing, like splitting and compressing files inside a tar archive. The second solution would be to implement a new functionality in Biobtree and save and decompress these files to local disk and make parallel access to decompressed files.

A further obstacle would have happened during the generate phase, since we would have obtained more files from update phase. The generate phase struggles to merge all these files and could cause much longer times of output generation. To address this obstacle, a new phase could be implemented that runs before the generate phase and merges files coming from the update phase.

## Limitations

Although duplicate values for each key are discarded during the update phase for each dataset, the generate phase could rarely produce duplicate values. The user needs to discard these duplicate records manually if these are created. Another limitation is when querying the special keywords, they need to be fully specified including all space characters.

## Future Work

The limitations can be addressed and new functionalities added in the future. For instance, different bioinformatics datasets like Ensembl [7] or ENA [8] can be integrated. Another feature would be sorting result values based on a certain criterion.

## Competing Interests

No competing interests were disclosed.

## Data (and Software) Availability

All source codes and binaries are availabe at https://www.github.com/tamerh/biobtree

## Grant Information

The author(s) declared that no grants were involved in supporting this work.

